# Functional Inputs to the Subgenual Cingulate Cortex Distinguish Neurotypical Individuals with Severe and Mild Depressive Problems: Granger Causality and Clustering Analyses

**DOI:** 10.1101/2025.06.25.661556

**Authors:** Huey-Ting Li, Yu Chen, Shefali Chaudhary, Jaime S. Ide, Chiang-Shan R. Li

## Abstract

Extant research has implicated functional connectivity of the subgenual anterior cingulate cortex (sgACC) in major depressive disorders or depressive traits in neurotypical populations. However, prior studies have not distinguished the inputs and outputs of the sgACC, and the “diagnostic” accuracy of these connectivity metrics remains elusive. Here, we analyzed data of 890 subjects (459 women, age 22 to 35) from the Human Connectome Project using Granger causality analyses (GCA) with the sgACC as the seed and 268 regions of interest from the Shen’s atlas as targets. Individual connectivities were assessed with an *F* test and group results were evaluated with a binomial test, both at a corrected threshold. We identified brain regions with significant input to and output from the sgACC. Clustering analyses of Granger causality input, but not Granger causality output or resting state connectivity features revealed distinct subject clusters, effectively distinguishing individuals with severe and mild depressive symptoms and those with comorbidities. Specifically, weaker projections from the fronto-parietal and orbitofrontal cortices, anterior insula, temporal cortices, and cerebellum to the sgACC characterized five clusters with low to high scores of depression as well as comorbid internalizing and externalizing problems. Machine learning using a logistic classifier with the significant “GCA-in” features and 5-fold cross-validation achieved 87% accuracy in distinguishing subject clusters, including those with high vs. low depression. These new findings specify the functional inputs and outputs of the sgACC and highlight an outsized role of sgACC inputs in distinguishing individuals with depressive and comorbid problems.

**Significance Statement:** Depression affects millions worldwide. Understanding the neural mechanisms would facilitate treatment of depression. In this study, we investigated how different parts of the brain send signals to and/or receive signals from a key brain region – subgenual cingulate cortex – implicated in the pathophysiology of depression. Using brain scans from nearly 900 adults, we found that weaker signals coming into this area, especially from regions involved in thinking and emotion, were linked to higher levels of depression and other behavioral problems such as anxiety and aggression. These findings help us better understand how regional communications in the brain relate to emotional well-being and may support new ways to identify and treat people with a depressive disorder.

## Introduction

The subgenual anterior cingulate cortex (sgACC), a key hub of the emotion-regulating limbic circuit (Disner et al., 2011), is widely implicated in the pathophysiology of mood disorders (Drevets et al., 2008; Hajek et al., 2008). Altered sgACC activity has been observed in individuals with major depressive disorder (MDD) (Gotlib et al., 2005; Matthews et al., 2008; Stuhrmann et al., 2011). Greater sgACC activation to sad emotional stimuli early during treatment predicted improvement in depression scores with 10 weeks of antidepressant regimen (Keedwell et al., 2010). In neurotypical populations, elevated sgACC responses to social exclusion predicted later depressive symptoms in adolescents (Masten et al., 2011). These findings highlight sgACC’s crucial role in emotion processing and sgACC dysfunction as a neural marker for depression and treatment response.

Deep brain stimulation (DBS) and transcranial magnetic stimulation (TMS) targeting the sgACC have shown promise in treating people with depression (Berlim et al., 2014). Chronic DBS of white matter tracts near the sgACC led to sustained remission for up to six months (Mayberg et al., 2005), with at least 50% reduction in depression scores (Merkl et al., 2016). DBS in the sgACC also improved empathic responses to negative stimuli and reduced depression severity in treatment-resistant MDD (Merkl et al., 2016). Repetitive TMS of the dorsolateral prefrontal cortex (dlPFC) modulated sgACC activity in depressed patients (Duprat et al., 2025), with treatment responders showing negative sgACC-dlPFC connectivity at rest (Fox et al., 2012). Further, baseline sgACC-frontoparietal connectivity predicted response to escitalopram over 12 weeks of treatment (Wang et al., 2024), highlighting sgACC functional connectivity as a biomarker for treatment efficacy.

Resting-state functional connectivity (rsFC) reveals brain’s functional architecture by quantifying regional correlations of spontaneous neural activity, independent of task demands (van den Heuvel and Pol, 2010). Numerous studies have highlighted altered rsFC in neuropsychiatric disorders, including depression and anxiety, reflecting intrinsic brain networks in link with cognitive dysfunction and emotional dysregulation (Mulders et al., 2015; Tang et al., 2018). Studies showed altered sgACC rsFC’s with frontoparietal, salience, limbic-striatal, and default mode networks (DMN) in depression (Zhou et al., 2024). Relative to healthy controls, MDD patients showed higher sgACC rsFC’s with the prefrontal cortex (PFC; Davey et al., 2012; Peng et al., 2021) and amygdala (Connolly et al., 2013; Ho et al., 2014) and lower rsFC’s with the posterior cingulate cortex (PCC), hippocampus, thalamus, and insula (Peng et al., 2021; Zhang et al., 2022). These connectivity patterns correlated with depression severity in both clinical (Davey et al., 2012; Ho et al., 2014; Peng et al., 2021; Zhang et al., 2022) and subclinical (Philippi et al., 2015; Chen et al., 2025) populations.

Whereas the rsFC describes correlated brain activities, it does not determine the direction of influence. Granger causality analysis (GCA), a statistical method for assessing directional relationships in time-series data, helps in distinguishing inputs from outputs (Seth et al., 2015). For instance, GCA demonstrated enhanced bottom-up connections from the thalamus to cortical and subcortical regions, along with reduced top-down connections to the thalamus, in people with MDD (Yang et al., 2023). On the other hand, despite extensive evidence implicating the sgACC in depression, no studies have distinguished the roles of sgACC inputs and outputs in depression.

To address this issue, we investigated Granger causal connectivities of 268 regions of interest (ROIs; Shen et al., 2013; Finn et al., 2015) with the sgACC in 890 neurotypical individuals curated from the Human Connectome Project. After identifying ROIs with significant sgACC inputs and outputs, we used connectivity strength (*F* values) in cluster analysis to identify groups with distinct connectivity features. We hypothesized that these features could differentiate individuals with varying depression and/or anxiety scores and potential comorbidities. Lastly, we applied machine learning to evaluate the accuracy at which these features predicted group membership. Our overall goal is to define sgACC functional inputs and outputs and characterize their potentially distinct roles in individual depression.

## Materials and Methods

### Dataset: Subjects and Assessment

The HCP Young Adult (HCP-YA) S1200 release contains clinical, behavioral, and 3T magnetic resonance (MR) imaging data of 1,206 young healthy adults without severe neurodevelopmental, neuropsychiatric, or neurologic disorders. In the current study, a total of 316 subjects were excluded due to missing resting-state fMRI (rsfMRI) or clinical data (*n*=110), excessive head movements and non-stationary time series (*n*=206, see details in *MRI Data Preprocessing and Granger Causality Analysis*). Therefore, the final cohort comprised 890 subjects (459 women, age 22 to 35). Participants completed the Achenbach Adult Self Report (ASR; Achenbach and Rescorla, 2003), where individual questions were rated on a 3-point Likert scale (0-Not True, 1-Somewhat or Sometimes True, and 2-Very True or Often True). We used the Diagnostic and Statistical Manual-oriented depression and anxiety raw scores in our analyses. The ASR also includes 8 syndromal scales, including anxious/depressed syndromes; withdrawn syndromes; somatic complaints; thought problems; inattention problems; aggression; rule-breaking behaviors; and intrusive problems, and allowed us to evaluate comorbidities of the subject clusters. For depression, anxiety, and all comorbid syndromes, higher scores indicate greater symptom severity.

### MRI Protocol

As described earlier (Chen et al., 2021; Chen et al., 2024), the details of the data collection procedures can be found in the HCP S1200 Release Reference Manual. All imaging data were acquired on a customized Siemens 3T Skyra with a standard 32-channel Siemens receiver head coil and a body transmission coil. T1-weighted high-resolution structural images were acquired using a 3D MPRAGE sequence with 0.7mm isotropic resolution (FOV=224mm, matrix=320, 256 sagittal slices, TR=2400ms, TE=2.14ms, TI=1000ms, FA=8°) and used to register rsfMRI data to a standard brain space. The rsfMRI data were collected in two sessions, using gradient-echo echo-planar imaging (EPI) with 2.0 mm isotropic resolution (FOV=208×180mm, matrix=104×90, 72 slices, TR=720ms, TE=33.1ms, FA=52°, multi-band factor=8). Within each session, oblique axial acquisitions alternated between phase encoding in a right-to-left direction in one run and left-to-right direction in the other run. Each run lasted 14.4 minutes (1200 frames).

### MRI Data Preprocessing and Granger Causality Analysis (GCA)

Preprocessing was applied to reduce spurious BOLD variances that were unlikely to reflect neuronal activity (Fox and Raichle, 2007; Zhang et al., 2012). The sources of spurious variance were removed by regressing out signals from the ventricular system, white matter, and whole brain, in addition to the 6 parameters obtained by rigid body head motion correction. Additionally, we applied a temporal band-pass filter (0.009Hz<*f*<0.08Hz) to the time course in order to obtain low-frequency fluctuations (Dong et al., 2023). Cordes et al. (2001) suggested that BOLD fluctuations below a frequency of 0.1Hz contribute to regionally specific BOLD correlations. Next, average time series were extracted for ROIs defined according to the Yale brain atlas of 268 regions covering the whole brain (Shen et al., 2013) as well as for the sgACC.

Granger causality between the sgACC and each of the 268 ROIs was computed using *F*-values to represent the causality strength (Goebel et al., 2003). With a TR of 0.72 s, we used model order 2 rather than higher orders to avoid model overfitting in GCA (Roebroeck et al., 2005). We called the influence of sgACC on other ROIs “*GCA*-out”, and the influence of ROIs on the sgACC “*GCA*-in.” For both *GCA*-in and *GCA*-out, the connectivity significance for each individual was assessed using the *F*-statistics of autoregressive modeling. Multiple comparisons across ROIs were evaluated at *p*<0.01, corrected for false discovery rate (FDR). Group results were evaluated with a binomial test of the number of subjects with versus without a significant input or output connection for each ROI, again at *p* < 0.01, Bonferroni corrected (Ide and Li, 2011).

In addition to *GCA*-in and *GCA*-out, we also computed the rsFC’s of the sgACC with the 268 ROIs, using the same preprocessed BOLD data and following a published pipeline (Chen et al., 2021; Zhang et al., 2021). Pearson’s correlations were computed pairwise, and *z*-transform was applied to the coefficients for one-sample *t*-test in group analysis. These rsFC’s served as features in a “control” set of clustering analysis, too.

### Clustering and Statistical Analyses

The connectivity metrics were divided into *GCA*-in and *GCA*-out features, each representing the *F*-in and *F*-out values, respectively, for significant ROIs, i.e., those with significant sgACC inputs or outputs. We applied *k*-means clustering to *GCA*-in and *GCA*-out features, respectively, and determined the optimum number of clusters (optimum*-k*) using the GAP statistics (Tibshirani et al., 2001). The GAP(*k*) statistics quantify the within-cluster dispersion of the actual data relative to randomly generated reference datasets (null distribution). The reference datasets are simulated in a way that preserves the original data (i.e., the overall shape in principal component space). The optimum-*k* was selected for the first *k* where the GAP statistic was not significantly smaller than the statistic for *k*+1, considering the standard deviations derived from the Monte-Carlo simulations. This is a more robust approach than simply selecting the maximum gap statistic because it considers the variability in GAP statistic estimates. The clusters represented distinct groups of subjects distinguished according to *GCA*-in or *GCA*-out metrics. In other words, subjects with similar GCA connectivity patterns were grouped together in a cluster. As a control, we also performed clustering analysis using the 268 sets of rsFC’s as features.

We next investigated if the subject clusters, with different GCA connectivity patterns, differed in demographic and ASR measures. We performed a one-way ANOVA, followed by post-hoc *t*-tests pairwise of the clusters, to examine whether the subject clusters differed in age and/or different levels of symptom severity. Differences in sex composition across the clusters were examined pairwise by chi-square tests.

To examine which GCA features were associated with depression severity, we ordered the cluster-labels according to the ASR scores for correlation with the neural features (i.e., *GCA*-in or -out F values). We ordered the cluster-labels by increasing value of depression scores, so that a significant negative correlation indicated that lower GCA connectivity (*F* value) of a particular ROI with the sgACC was associated with higher symptom severity of depression, and vice versa. To assess the significance of the correlations, we employed a permutation test. This non-parametric method provides a robust way to estimate empirical *p*-values without making strong assumptions about the data distribution, with the following procedures: i) *calculate correlations*: we first calculated the Pearson’s correlation coefficients between each connectivity feature and the cluster label; ii) *generate null distribution*: to create a null distribution, we randomly permuted the cluster labels while keeping the feature values fixed. This removed the relationships between the features and clusters, simulating a scenario where there was no association. For each permutation, we calculated the correlation coefficients between the features and shuffled labels. This process was repeated 10,000 times to generate a distribution of correlation coefficients under the null hypothesis of no association; and iii) *estimate empirical p-values*: for each feature, we compared the observed correlation coefficient to the null distribution. The empirical *p*-value was calculated as the proportion of permuted correlation coefficients that were greater than or equal to the absolute value of the observed correlation coefficient. This provides an estimate of the probability of observing a correlation as extreme as the observed one if there were no true association between the feature and cluster labels.

### Predicting Depression using Multi-label Classification

To assess whether and how well the GCA features could predict levels of depression, we used the ordered cluster-labels as class-labels for supervised learning. This approach has the benefit of filtering out variance present in the clinical measures. We employed two methods – multinomial logistic and naïve Bayes modeling – in the multi-label classification.

The analysis followed a two-stage pipeline: training and testing (**Supplementary Figure S1**). The classifier was trained on a portion of the clustered data, using 5-fold cross-validation to assess their performance on unseen data. This cross-validation procedure allowed us to evaluate the performance of the classifiers. To quantify classification performance, we used accuracy, balanced accuracy, and macro-averaged F1-score as performance metrics. Accuracy was the overall proportion of correctly classified instances. Balanced accuracy was the average recall for each class. Macro-averaged F1-score was the average of the F1-score for each class, where the F1-score was the harmonic mean of precision and recall. These metrics provided a comprehensive evaluation of the classifier’s performance across all classes, especially in cases where the classes may have different sizes or importance. The entire pipeline was implemented using Python (scikit-learn==1.0.2).

## Results

### Granger causality and rsFC maps of the sgACC

Figure 1A shows the ROIs with significant inputs and outputs of the sgACC as shown by GCA. The frontopolar cortex, bilateral superior/middle/inferior gyri, ventromedial PFC, inferior temporal cortex (ITC), and inferior parietal cortex, precuneus, anterior insula (AI) as well as the cerebellum provide significant inputs to the sgACC.

**Figure 1.**
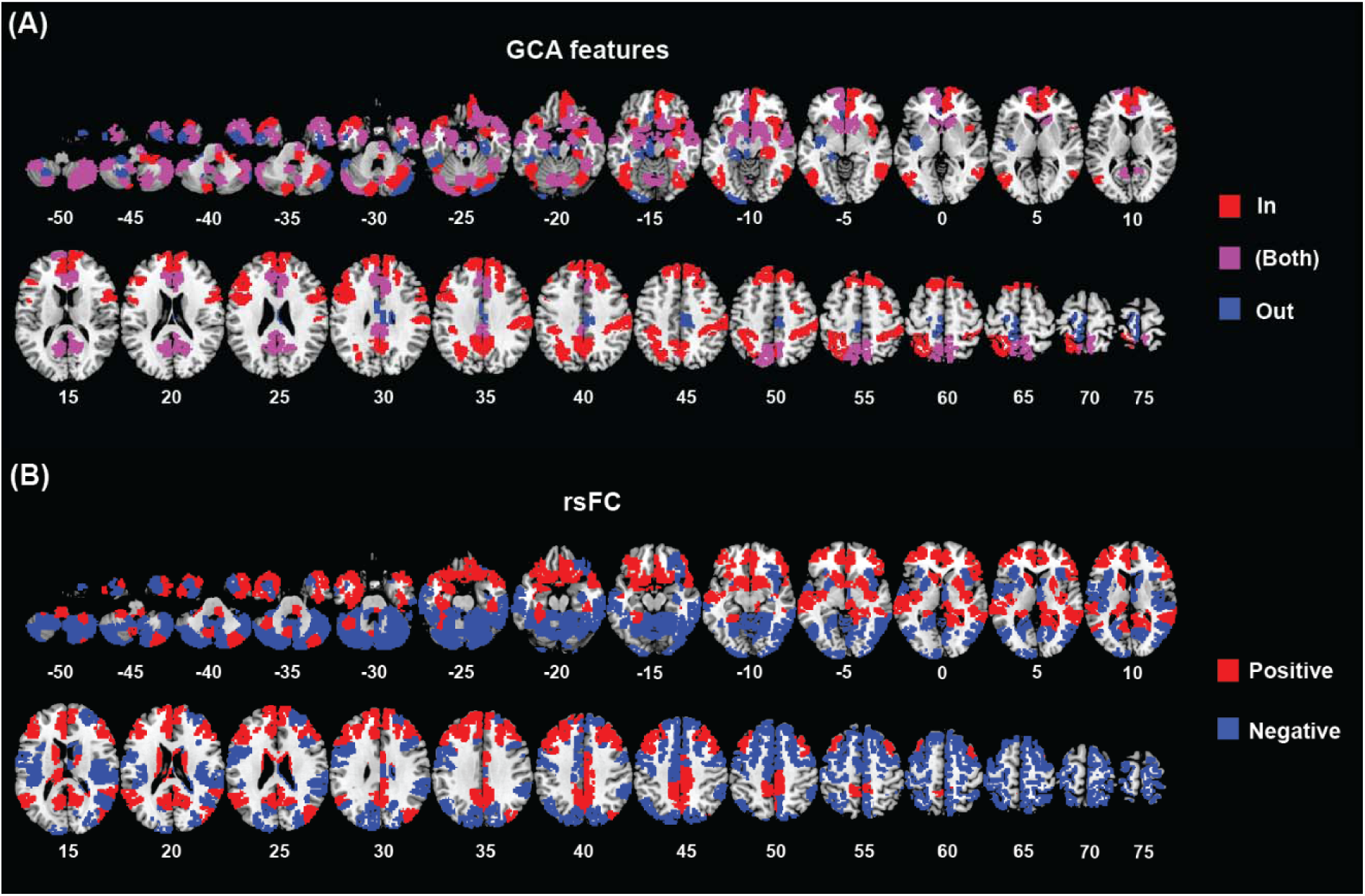
Granger causality and rsFC maps of the sgACC. **(A)** Regions with *GCA*-in and *GCA*-out connectivity with the sgACC. GCA metrics are thresholded at *p*<0.01, corrected for false discovery rate (FDR), for individual subjects, and binomial test was used for group analyses with *p*<0.01, Bonferroni corrected for multiple comparisons across regions of interest (ROIs). Red, blue, and violet colors indicate regions with inputs to, outputs from, and both inputs to and outputs from the sgACC, respectively. **(B)** Regions functionally connected to the sgACC revealed by resting state connectivity analysis. The map showed one sample *t*-test result of *z*-transformed correlation coefficients at *p*<0.001, Bonferroni corrected for multiple comparisons across ROIs. Red and blue colors indicate regions in positive and negative correlation with the sgACC, respectively.

The medial motor cortex, premotor cortex, including the supplementary motor area, mid-cingulate cortex (MCC), brain stem, cerebellum and part of bilateral ITC and left occipital cortex received significant projections from the sgACC.

Some of these brain regions both provided inputs to and received outputs from the sgACC, including the rostral ACC (rACC), PCC, precuneus, bilateral caudate head, hippocampus/parahippocampal gyri, posterior orbitofrontal cortex (OFC), and cerebellum, as well as the anterior part of the right ITC.

One-sample *t* test of the sgACC rsFC’s showed extensive positive and negative connectivities encompassing the frontal, parietal, temporal, and occipital cortices, insula, cerebellum, and subcortical regions (Figure 1B).

### Cluster differences in clinical features

#### Clustering with significant GCA-in features

We employed *GCA*-in features (*F* values) of all 84 ROIs that were significant at *p*<0.01 for both individual and group level analyses in *k*-means clustering. In Figure 2A, we show the GAP statistics for different numbers of clusters going from 1 to 15. The optimal clustering was selected at *k*=5 for the *GCA*-in features (see Methods). The mean and standard deviation of demographic and ASR measures of these five clusters are shown in **Supplementary Table S1A**, with cluster labels ordered according to increasing levels of average depression scores. Pair-wise cluster differences in demographic and ASR measures are summarized in **Supplementary Table S1B**.

**Figure 2.**
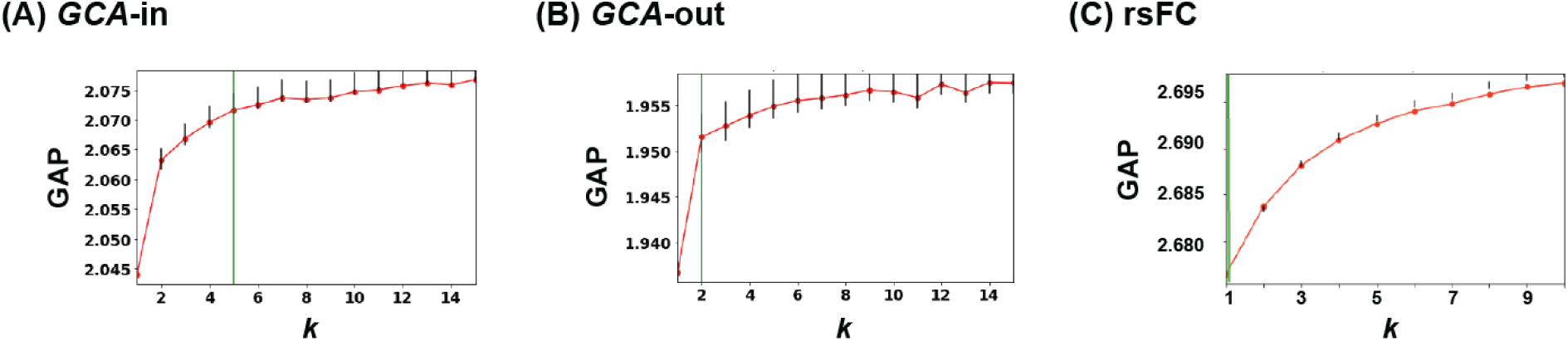
GAP statistics of clustering based on **(A)** *GCA*-in, **(B)** *GCA*-out, and **(C)** rsFC features. Y-axis represents the GAP statistic – the ratio between within-cluster dispersion of the actual data as compared to random data. A higher GAP value suggests better clustering. The green line represents the selected optimum number of clusters (*k*=5, *k*=2, and *k*=13 for *GCA*-in, *GCA*-out, and rsFC, respectively). The error bars represent the standard deviation.

As shown in Figure 3, Cluster C1 showed the lowest scores of depression (**3A**) and anxious/depressed syndromes (**3C**), significantly different from those of C3, C4, and C5. Cluster C1 vs. C4 and C5, as well as Cluster C2 vs. Cluster C4 showed significant lower anxiety scores (**3B**). Cluster C1 also showed the lowest score of somatic complaints, significantly different from that of C2 and C5 (**3E**). Relative to C5, Cluster C1 showed significant lower scores of inattention problems (**3G**). Cluster C1 showed lower aggression score than C4 and C5 (**3H**). Cluster C4 showed the lowest score of rule-breaking behaviors, which were significantly different from the score of C5, the highest of all (**3I**). In addition, the sex composition of Cluster C4 (female-to-male ratio=1.81:1) differed significantly from that of C1 (0.77:1), C2 (0.78:1), and C5 (1:1) (**3K**). There were no significant differences between clusters in withdrawn syndromes (**3D**), thought (**3F**), or intrusive problems (**3J**).

**Figure 3.**
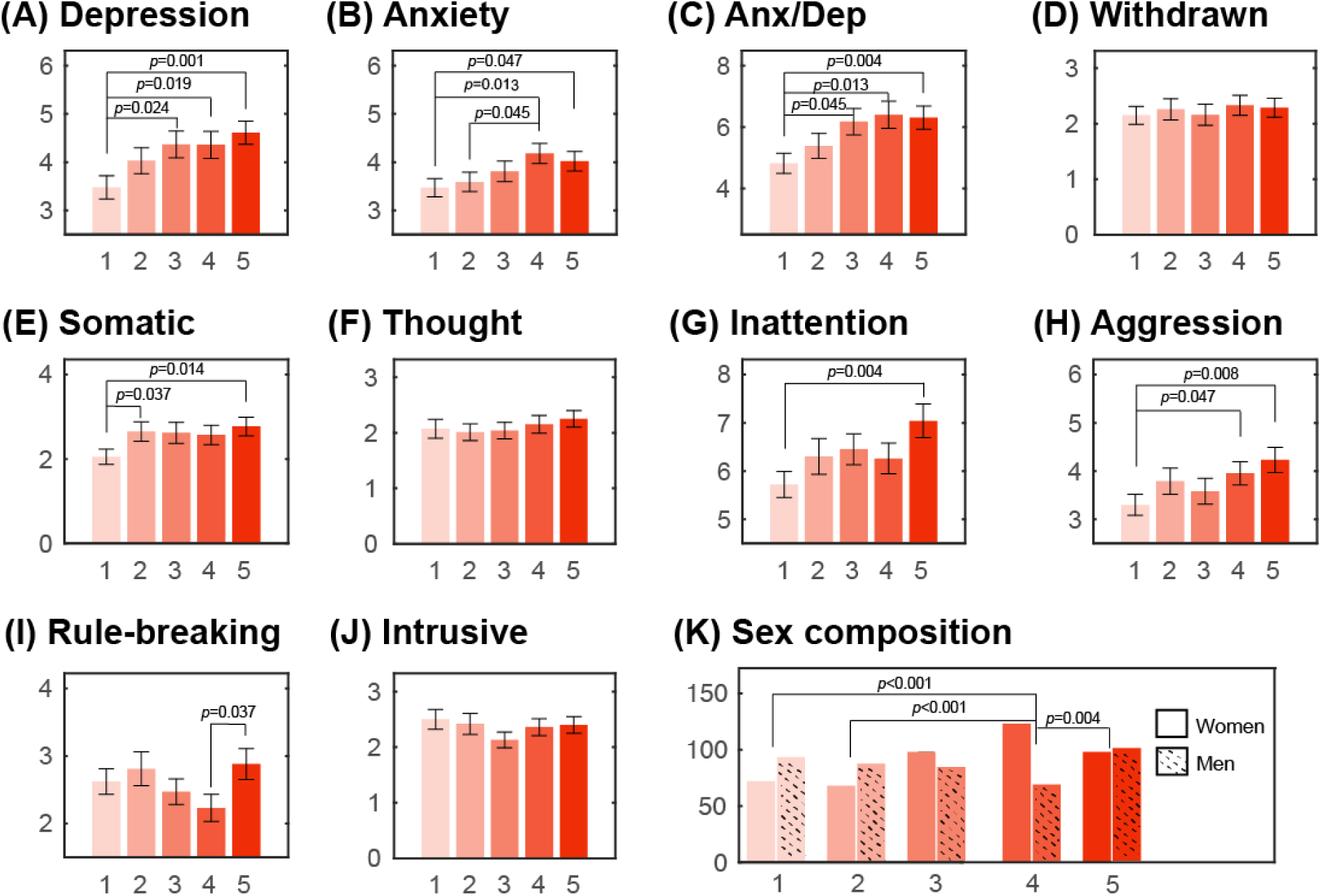
Pair-wise comparison in ASR measures between clusters based on significant GCA-in features. **(A)** Depression; **(B)** Anxiety; **(C)** Anxious/Depressed syndromes; **(D)** Withdrawn syndromes; **(E)** Somatic complaints; **(F)** Thought problems; **(G)** Inattention problems; **(H)** Aggression; **(I)** Rule-breaking behaviors; **(J)** Intrusive problems; **(K)** Sex composition. X- and Y-axis represent the clusters C1 to C5 based on significant GCA-in features and ASR raw scores, respectively. Pair-wise comparison with *p*’s<0.05 was also presented. ASR: Adult Self Report.

To summarize the clinical characteristics of these subject clusters, C1 and C5 were characterized by the lowest and highest levels across different domains of internalizing and externalizing symptoms.

#### Clustering with significant GCA-out features

In Figure 2B, we show the GAP statistics for the 64 significant *GCA*-out features. The optimal clustering was selected at *k*=2 for the *GCA*-out features. There were no statistically significant differences of clinical or demographic variables between the two clusters (*p*’s≥0.130), except for sex (*χ^2^*=6.54, *p*=0.010), with more men in Cluster 1 (53% of a total of 367) and more women in Cluster 2 (55% of a total of 523).

#### Clustering with rsFC features

We applied the same clustering procedures to rsFC metrics (Pearson’s correlations) and obtained *k*=13 as the optimum number of clusters (Figure 2C). However, inspection of the clusters revealed issues that suggested invalidity of the results.

As shown in the Supplement, we expanded the range and computed the GAP statistics for rsFC clustering up to *k*=15, as well as examined the data distribution (**Supplementary Figure S2A**) and within-cluster sum of squared distances (Wks) to further investigate the existence of a larger number of clusters. According to the 1-SE criterion, we obtained *k*=13 as the optimum number of clusters; however, inspection of the cluster distribution showed absence of well-defined clusters (**Supplementary Figure S2B** and **2C**), indicating the existence of a non-monotone behavior that violated one of the key assumptions of GAP statistics (Tibshirani et al., 2001). Further, the decrease of the Wks could be explained by the creation of smaller subclusters, including those with only a single metric (**Supplementary Figure S2D**), again suggesting overestimation of the number of clusters (Lange et al., 2004). We also estimated the optimal number of clusters using Silhouette scores and confirmed the results. Density-Based Spatial Clustering of Applications with Noise, another algorithm that groups data points based on their proximity to each other, likewise failed to show multiple clusters. Therefore, we concluded that rsFC’s as features did not yield significant clustering, i.e., *k*=1.

Finally, following the clustering results with GCA metrics as features, we arbitrarily selected *k*=4 and *k*=5 for clustering with rsFC as features and examined whether the four or five clusters may show any group differences in depression scores. The results were negative (*t’s*≤1.90; *p’s*≥0.050; pair-wise *t* tests).

### Depression-related GCA-in features: identification and prediction

The depression score increased from GCA-in Cluster C1 to C5, with statistical differences observed between C1 and C3-5. Therefore, we examined which *GCA*-in feature (GC strength or *F* value) were significantly associated with depression by computing the correlation between the *F*-in values and the ordered cluster-labels. The brain regions with significant correlations as evaluated with permutation tests, are shown in Figure 4 and listed in **Supplementary Table S2**. Note that these correlations were all negative, indicating that lower GC strength was associated with higher severity of depression symptoms. The critical *GCA*-in features included projections to the sgACC from the frontal, parietal, temporal, and occipital areas, specifically, bilateral superior frontal and middle temporal gyri, left posterior orbital gyrus and superior parietal lobule, right medial frontal cortex and precentral gyrus.

**Figure 4.**
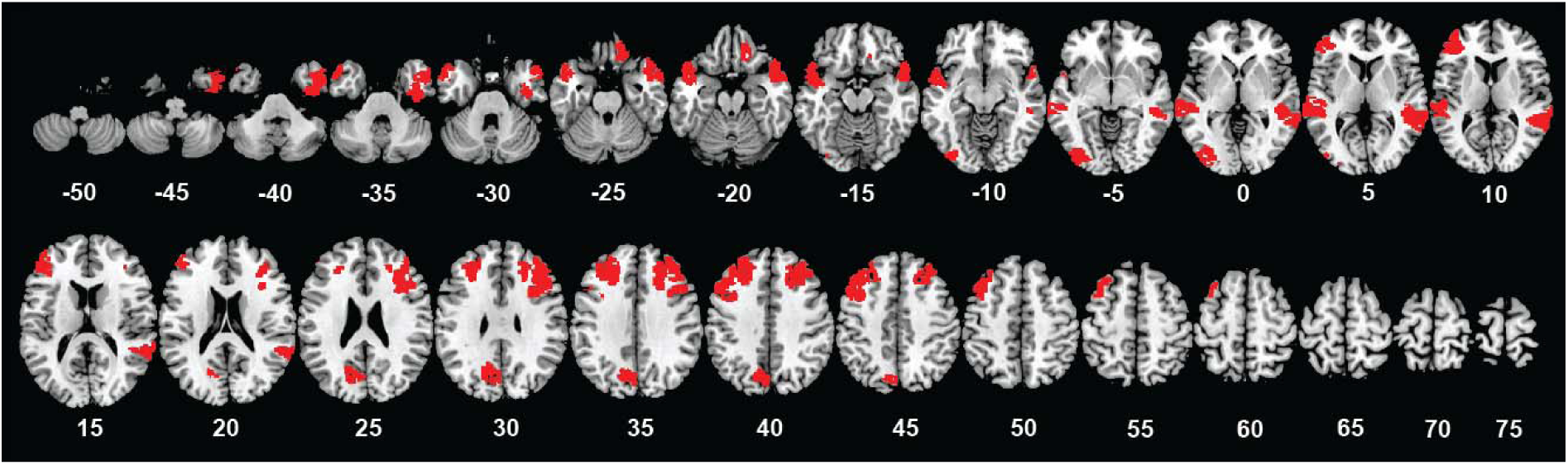
Brain regions with significant correlation between significant *GCA*-in features of the sgACC and cluster-labels ordered by depression. Results were evaluated at *p’s*<0.001, Bonferroni corrected. All *GCA*-in features showed negative correlation with cluster labels in increasing severity of depression. We illustrated these input ROIs in red to be consistent with Figure 1.

We used *GCA*-in features to predict the clusters with significant differences in depression. The multinomial logistic model showed an accuracy of 86.7%, a balanced accuracy of 86.5%, and an F1 score of 0.865. The Naïve Bayes model showed an accuracy of 80.4%, a balanced accuracy of 79.9%, and an F1 score of 0.800.

## Discussion

Significant Granger causal inputs to the sgACC enabled clustering of individuals with varying severity of depression and comorbid problems. Connectivity features of multiple cortical regions best captured depression and predicted its severity with 86.7% accuracy in a logistic model. Notably, the same analytics applied to Granger causal outputs or to rsFC’s failed to identify depression-related subject clusters. These findings suggest that functional connections influencing sgACC resting-state BOLD fluctuations may underlie individual differences in depression severity in neurotypical young adults.

### Functional connectivities of the sgACC: rsFC analysis and GCA

The sgACC rsFC’s showed both positive and negative connections with frontal, parietal, temporal, and occipital cortices, as well as the insula, cerebellum, and subcortical regions, in alignment with prior findings (Margulies et al., 2007; Wang et al., 2014). This suggests that the sgACC partakes in large-scale functional brain networks and, by integrating information across multiple cortical and subcortical regions (Barrett and Satpute, 2013), may support emotional regulation and other cognitive processes critical to the pathophysiology of mood disorders (Drevets et al., 2008).

Adding to studies of rsFC, we identified an extensive array of brain regions that were either unidirectional recipients (*GCA*-in) or sources (*GCA*-out) or of bidirectional influences of the sgACC. Regions providing inputs to the sgACC encompassed the fronto-parietal network, frontopolar cortex, and DMN (including the precuneus), ITC, and AI. This network of inputs likely regulates sgACC activity. For example, the frontopolar cortex (Adamczyk et al., 2025) and lateral PFC (Ochsner and Gross, 2005; Hamilton et al., 2012) support emotion regulation, with direct anatomical connections (Hogeveen et al., 2022), may regulate sgACC activities and emotional responses. Anatomical tract-tracing in nonhuman primates showed that dlPFC fibers extend ventrally and medially, merging with cingulate bundles before terminating in the sgACC (Heilbronner and Haber, 2014). These prefrontal cortical connections support sgACC’s role in cognitive-affective processing.

The precuneus, a key DMN component, may influence sgACC activity in self-referential negative emotions (Whitfield-Gabrieli and Ford, 2012; Nejad et al., 2013). The sgACC-precuneus hyperconnectivity has been associated with rumination and difficulty disengaging from negative thoughts (Sheline et al., 2010; Marchetti et al., 2012). Brain stimulation and psychotherapy alleviated depression by disrupting this hyperconnectivity (Liston et al., 2014).

Involved in interoception, emotional awareness, and salience processing (Craig, 2009; Menon and Uddin, 2010; Gu et al., 2013; Wang et al., 2019), the AI functions as a key limbic input region (Medford and Critchley, 2010). GCA showed positive causal outflow from the right AI to the pgACC in both MDD patients and healthy controls (Iwabuchi et al., 2014). Our findings add to this literature and support a potential role of AI-sgACC connectivity in the regulation of interoceptive, salience and emotion responses and in the pathophysiology of depression.

In contrast, the sgACC predominantly sent outputs to medial motor cortical structures, the cerebellum, MCC, pons, and left posterior insula. The fronto-pontine-cerebellar circuits may support emotional expression (Schmahmann and Pandya, 1997; Paus, 2001; Stoodley and Schmahmann, 2009; Holroyd and Yeung, 2012), and sgACC outputs to the posterior insula modulate autonomic and visceral states of emotions (Critchley et al., 2004; Craig, 2009). As a core region for autonomic regulation, the sgACC influences heart rate variability and stress responses (Thayer et al., 2012) and, with its projections to the posterior insula, may modulate bodily states during emotional processing in anxiety and depression (Wiebking et al., 2014).

Additionally, a wide array of brain regions showed bidirectional connectivity with the sgACC, including the rostral ACC, PCC, dorsal parietal cortex, ventral striatum, hippocampus, amygdala, and right insula. These connections are linked functionally to emotional regulation, attention to emotional stimuli, and reward sensitivity (Corbetta and Shulman, 2002; Haber and Knutson, 2010). Dysfunctional sgACC-ventral striatum connectivity was associated with anhedonia (Russo and Nestler, 2013) and sgACC-hippocampal hyperconnectivity was associated with excessive emotional recall in depression (Neumeister et al., 2006). Together, these findings are in broad agreement with the roles of sgACC network (Zhang et al., 2017; Chen et al., 2020) and extend the literature by specifying the directional interactions of the sgACC and how these directional interactions may play out in depression.

### Input features of the sgACC predict individual variation in depression

Indeed, by leveraging *GCA*-in features in multi-label classification, we successfully predicted individual variations in depression problems. Notably, classification based exclusively on *GCA*-in, but not *GCA*-out, features effectively clustered participants by depression levels. This suggests that incoming influences on the sgACC, rather than its outbound signaling, help in distinguishing depression severity in this neurotypical population. Disruptions in how the sgACC processes inputs from the emotion circuit contributes significantly to depressive symptoms (Disner et al., 2011). Prior studies of GCA reported that medial PFC and hippocampal activity predicted higher ventral ACC activity in people with MDD (Hamilton et al., 2011). Additionally, connectivity from the OFC to the ACC and occipital regions was associated with depression scores in patients with MDD (Gao et al., 2016).

We found significant associations between the ordered cluster-labels of depression and Granger causal connectivity strength of these inputs to the sgACC, including those from bilateral frontal, temporal, occipital regions, left MCC, and precuneus. Earlier research showed that higher dlPFC activity led to reduced ventral ACC, PCC, dorsomedial PFC (dmPFC), and ventral striatum activation in MDD (Hamilton et al., 2011). In particular, sgACC-amygdala dysconnectivity was associated with heightened emotional reactivity (Johnstone et al., 2007) and emotion regulation enhanced dmPFC-amygdala connectivity via sgACC pathways (Scharnowski et al., 2020). These findings are consistent with labeled neurons in the amygdala following sgACC injections in neuronal tracing studies of macaque monkeys (Kim et al., 2018). While limited research has explored sgACC Granger causal connectivity, Zhou and colleagues reported directional influences of the sgACC and pregenual ACC (pgACC) on the right amygdala during emotional face processing in healthy adults (Zhou et al., 2011). Further, a study using dynamic causal modeling found weaker connectivity from the amygdala to the sgACC in adolescents with MDD (Musgrove et al., 2015). Here, we observed reciprocal influences between bilateral amygdala and sgACC (GCA-in and GCA-out both significant). However, amygdala-sgACC connections were not amongst those that distinguished cluster levels of depression. Thus, it remains to be seen whether Granger causality connectivities between the sgACC and amygdala may only be compromised in individuals with a clinical diagnosis of depression.

### Cluster differences in depression and comorbid problems

Clustering based on *GCA*-in *F* values of these sgACC-connected ROIs identified five clusters of individuals with distinct affective and behavioral profiles. The sgACC input-based clustering revealed clinically meaningful phenotypes: Cluster C1 exhibited the lowest levels of depression, anxiety, somatic complaints, and inattention problem scores, whereas C5 showed the highest symptom burden across both internalizing and externalizing domains. These findings align with prior evidence linking structural atrophy and altered functional connectivity of the sgACC with anxiety (Hakamata et al., 2020), somatic symptoms (Yu et al., 2023; Chen et al., 2025), depression-related attentional deficits (Fales et al., 2008; Grahek et al., 2018), and aggressive behaviors (Siever, 2008). This dimensional grouping, based on functional connectivity features, supports the utility of sgACC-directed input as a neurobiologically-informed marker of clinical heterogeneity in depression and comorbid problems. Notably, machine learning showed that these sgACC markers are highly accurate (>85% accuracy) in distinguishing cluster labels, outperforming resting-state connectivity measures reported in previous studies (Craddock et al., 2009; Wang et al., 2021).

### Limitations and conclusions

This study has limitations, including the lack of formal depression diagnoses in the HCP sample and insufficient sample size for investigations of sex differences. Despite these limitations, our findings identify directional sgACC connections that may be important to mood and behavioral regulation. Multiple regional inputs to the sgACC predicted individual differences in depression and comorbid clinical features. Confirmation and refining of these findings in clinical populations is warranted and will help us better understand and manage depressive disorders.

To conclude, our findings support the sgACC’s role in mood regulation by integrating network inputs (Disner et al., 2011). Altered sgACC inputs are associated with more severe depression and comorbidities in neurotypical populations.

## Acknowledgements

This study is supported by NIH grants AG072893 and DA051922. The NIH is otherwise not responsible for the conceptualization of the study, data collection and analysis, or in the decision in publishing the results. The HCP data are provided by the WU-Minn Consortium (Principal Investigators: David Van Essen and Kamil Ugurbil; 1U54MH091657) funded by the 16 NIH Institutes and Centers that support the NIH Blueprint for Neuroscience Research; and by the McDonnell Center for Systems Neuroscience at Washington University.

The authors declare no competing financial interests.

