## Supplementary Materials for "Functional Inputs to the Subgenual Cingulate Cortex Distinguish Neurotypical Individuals with Severe and Mild Depressive Problems: Granger Causality and Clustering Analyses"

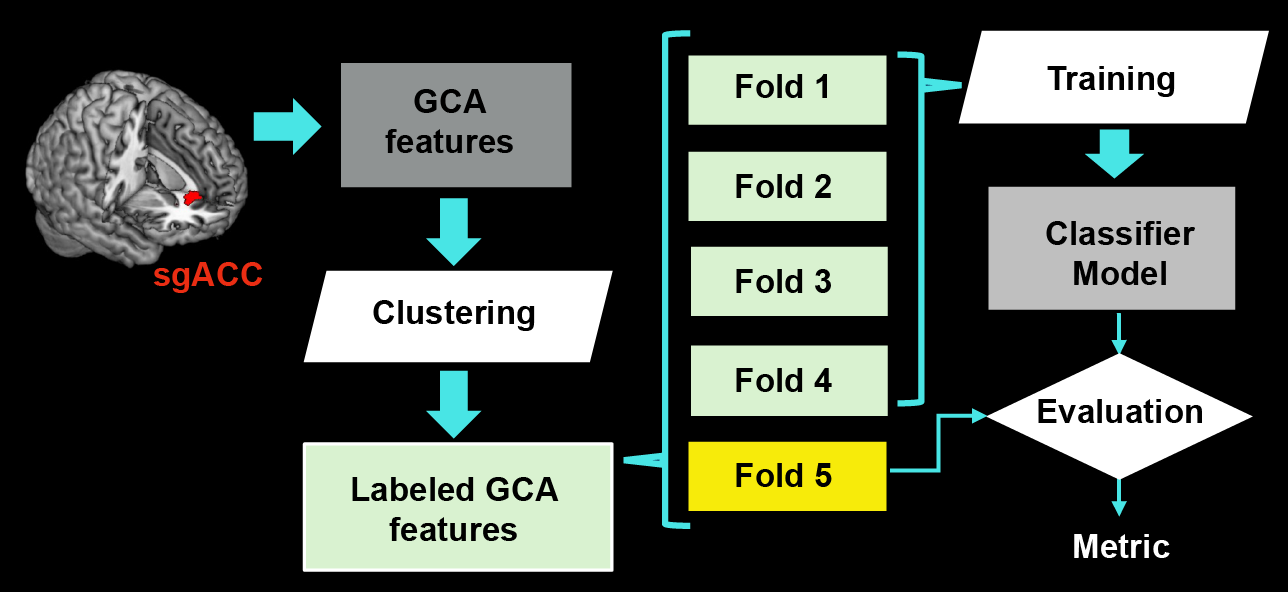

**Supplementary Figure S1.** Analytic pipeline for feature extraction, clustering, training, and testing of the accuracy of connectivity features in distinguishing subject clusters. The current diagram highlights the testing of fold-5, but all folds were tested to compute the metric mean and standard deviation in data analyses.

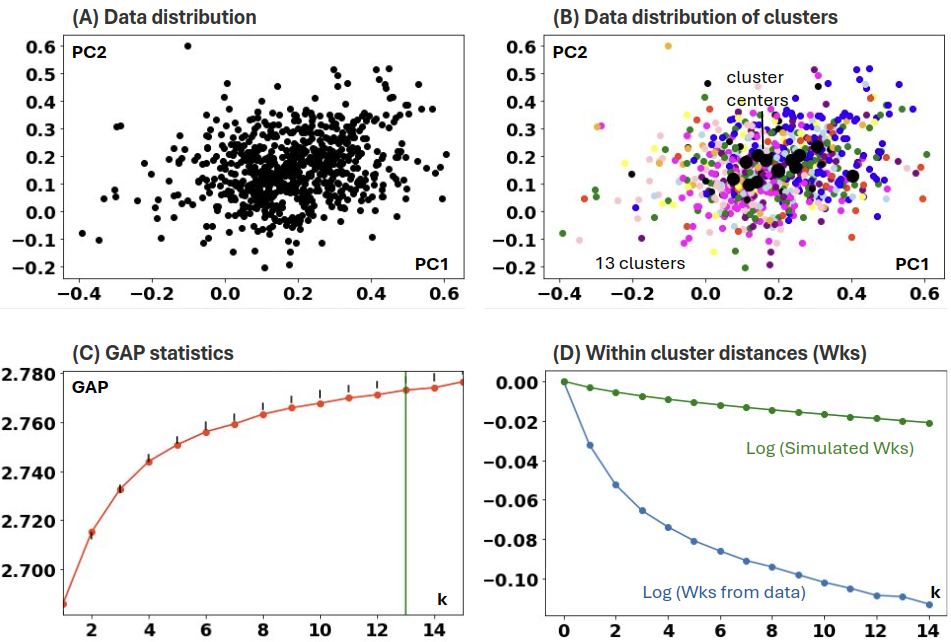

**Supplementary Figure S2.** Clustering of rsFC dataset. **(A)** Data distributions of the first two principal components (PC2) of the rsFC feature dataset; **(B)** Data distributions of the first two principal components (PC2) colored by the different clusters; **(C)** GAP statistics for different number of clusters; and **(D)** Comparison of within clusters sum of square distances (Wks) for rsFC features and simulated dataset.

**Supplementary Table S1A.** Clinical and demographic variables (mean±SD) of each subject cluster based on significant *GCA*-in features.

| **Variable** | **Cluster 1**  **(n = 165; 72 women)** | | **Cluster 2**  **(n = 155; 68 women)** | | **Cluster 3**  **(n = 181; 98 women)** | | **Cluster 4**  **(n = 191; 123 women)** | | **Cluster 5**  **(n = 198; 98 women)** | |
| --- | --- | --- | --- | --- | --- | --- | --- | --- | --- | --- |
|  | **mean** | **SD** | **mean** | **SD** | **mean** | **SD** | **mean** | **SD** | **mean** | **SD** |
| Depression | 3.480 | 3.049 | 4.029 | 3.329 | 4.326 | 3.794 | 4.355 | 3.839 | 4.611 | 3.430 |
| Anxiety | 3.466 | 2.408 | 3.586 | 2.446 | 3.807 | 2.835 | 4.177 | 2.902 | 4.015 | 2.759 |
| Age | 28.679 | 3.606 | 28.755 | 3.630 | 28.829 | 3.787 | 28.560 | 3.640 | 28.672 | 3.837 |
| Anxious/Depressed | 4.824 | 4.232 | 5.393 | 5.054 | 6.177 | 5.801 | 6.397 | 6.112 | 6.313 | 5.327 |
| Withdrawn | 2.147 | 2.084 | 2.262 | 2.311 | 2.155 | 2.589 | 2.329 | 2.481 | 2.293 | 2.352 |
| Somatic complaint | 2.046 | 2.263 | 2.646 | 2.823 | 2.624 | 3.296 | 2.569 | 3.161 | 2.768 | 3.107 |
| Thought problem | 2.074 | 2.122 | 2.009 | 1.913 | 2.044 | 2.043 | 2.149 | 2.209 | 2.247 | 2.138 |
| Inattention problem | 5.717 | 3.516 | 6.295 | 4.615 | 6.453 | 4.314 | 6.262 | 4.356 | 7.035 | 4.867 |
| Aggression | 3.296 | 2.790 | 3.785 | 3.339 | 3.575 | 3.649 | 3.945 | 3.274 | 4.227 | 3.707 |
| Rule-breaking behaviors | 2.622 | 2.423 | 2.808 | 3.166 | 2.470 | 2.597 | 2.229 | 2.710 | 2.884 | 3.226 |
| Intrusive problem | 2.500 | 2.247 | 2.419 | 2.390 | 2.127 | 1.869 | 2.358 | 2.116 | 2.399 | 2.157 |

**Supplementary Table S1B.** Differences in clinical and demographic variables pairwise amongst *GCA*-in subject clusters.

| **Variable** | | **C1 v. C2** | **C1 v. C3** | **C1 v. C4** | **C1 v. C5** | **C2 v. C3** | **C2 v. C4** | **C2 v. C5** | **C3 v. C4** | **C3 v. C5** | **C4 v. C5** |
| --- | --- | --- | --- | --- | --- | --- | --- | --- | --- | --- | --- |
| Depression | *t* | -1.534 | -2.265 | -2.347 | -3.280 | -0.754 | -0.830 | -1.598 | -0.072 | -0.766 | -0.693 |
|  | *p* | 0.126 | **0.024** | **0.019** | **0.001** | 0.451 | 0.407 | 0.111 | 0.942 | 0.444 | 0.488 |
| Anxiety | *t* | -0.441 | -1.194 | -2.484 | -1.993 | -0.754 | -2.013 | -1.518 | -1.241 | -0.724 | 0.563 |
|  | *p* | 0.660 | 0.233 | **0.013** | **0.047** | 0.452 | **0.045** | 0.130 | 0.215 | 0.470 | 0.574 |
| Age | *t* | -0.188 | -0.376 | 0.308 | 0.018 | -0.182 | 0.495 | 0.207 | 0.697 | 0.400 | -0.294 |
|  | *p* | 0.851 | 0.707 | 0.758 | 0.986 | 0.856 | 0.621 | 0.836 | 0.486 | 0.689 | 0.769 |
| Sex (W/M) | *χ^2^* | 0.000 | 3.404 | 14.577 | 1.016 | 3.126 | 13.761 | 0.890 | 3.638 | 0.643 | 8.203 |
|  | *p* | 1.000 | 0.065 | **<0.001** | 0.313 | 0.077 | **<0.001** | 0.346 | 0.056 | 0.423 | **0.004** |
| Anxious/Depressed | *t* | -1.091 | -2.451 | -2.772 | -2.899 | -1.306 | -1.636 | -1.643 | -0.355 | -0.238 | 0.144 |
|  | *p* | 0.276 | **0.015** | **0.006** | **0.004** | 0.192 | 0.103 | 0.101 | 0.723 | 0.812 | 0.886 |
| Withdrawn | *t* | -0.464 | -0.030 | -0.740 | -0.618 | 0.395 | -0.258 | -0.125 | -0.661 | -0.543 | 0.147 |
|  | *p* | 0.643 | 0.976 | 0.460 | 0.537 | 0.693 | 0.797 | 0.901 | 0.509 | 0.587 | 0.884 |
| Somatic complaint | *t* | -2.098 | -1.879 | -1.766 | -2.478 | 0.065 | 0.235 | -0.378 | 0.163 | -0.435 | -0.622 |
|  | *p* | **0.037** | 0.061 | 0.078 | **0.014** | 0.948 | 0.814 | 0.706 | 0.870 | 0.664 | 0.534 |
| Thought problem | *t* | 0.287 | 0.132 | -0.327 | -0.771 | -0.163 | -0.623 | -1.087 | -0.475 | -0.942 | -0.444 |
|  | *p* | 0.775 | 0.895 | 0.744 | 0.441 | 0.871 | 0.534 | 0.278 | 0.635 | 0.347 | 0.657 |
| Inattention problem | *t* | -1.260 | -1.724 | -1.282 | -2.896 | -0.323 | 0.067 | -1.446 | 0.423 | -1.225 | -1.644 |
|  | *p* | 0.209 | 0.086 | 0.201 | **0.004** | 0.747 | 0.946 | 0.149 | 0.673 | 0.221 | 0.101 |
| Aggression | *t* | -1.421 | -0.789 | -1.991 | -2.652 | 0.547 | -0.447 | -1.158 | -1.030 | -1.720 | -0.792 |
|  | *p* | 0.156 | 0.430 | **0.047** | **0.008** | 0.585 | 0.655 | 0.248 | 0.304 | 0.086 | 0.429 |
| Rule-breaking behaviors | *t* | -0.590 | 0.560 | 1.427 | -0.858 | 1.072 | 1.825 | -0.221 | 0.870 | -1.365 | -2.157 |
|  | *p* | 0.555 | 0.576 | 0.155 | 0.391 | 0.285 | 0.069 | 0.825 | 0.385 | 0.173 | **0.032** |
| Intrusive problem | *t* | 0.312 | 1.677 | 0.612 | 0.433 | 1.250 | 0.250 | 0.080 | -1.109 | -1.303 | -0.190 |
|  | *p* | 0.755 | 0.094 | 0.541 | 0.665 | 0.212 | 0.802 | 0.936 | 0.268 | 0.194 | 0.849 |

*Note:* A chi-square test was used to test sex composition, and two-sample *t* tests were used otherwise. Bold *p* values<0.05.

**Supplementary Table S2.** Significant depression-related regions with connectivity Granger influencing on the sgACC (only GCA significant regions)

| **Region** | ***r*** | ***p*** |
| --- | --- | --- |
| Left superior frontal gyrus | -0.358 | 1.19E-06 |
| Left superior frontal gyrus | -0.248 | 1.19E-06 |
| Left posterior orbital gyrus | -0.197 | 1.19E-06 |
| Left superior parietal lobule | -0.163 | 1.19E-06 |
| Left temporal pole | -0.181 | 1.19E-06 |
| Left temporal pole | -0.173 | 1.19E-06 |
| Left middle temporal gyrus | -0.275 | 1.19E-06 |
| Left middle temporal gyrus | -0.169 | 1.19E-06 |
| Right middle temporal gyrus | -0.230 | 1.19E-06 |
| Right temporal pole | -0.301 | 1.19E-06 |
| Right middle temporal gyrus | -0.214 | 1.19E-06 |
| Right temporal pole | -0.233 | 1.19E-06 |
| Right temporal pole | -0.254 | 1.19E-06 |
| Right superior frontal gyrus | -0.249 | 1.19E-06 |
| Right superior frontal gyrus medial segment | -0.267 | 1.19E-06 |
| Right medial frontal cortex | -0.164 | 1.19E-06 |
| Right precentral gyrus | -0.207 | 1.19E-06 |

*Note:* Permutation tests were evaluated at *p*<0.001, Bonferroni corrected.
